# Therapeutic efficacy of CT-P59 against P.1 variant of SARS-CoV-2

**DOI:** 10.1101/2021.07.08.451696

**Authors:** Dong-Kyun Ryu, Bobin Kang, Sun-Je Woo, Min-Ho Lee, Aloys SL Tijsma, Hanmi Noh, Jong-In Kim, Ji-Min Seo, Cheolmin Kim, Minsoo Kim, Eunji Yang, Gippeum Lim, Seong-Gyu Kim, Su-Kyeong Eo, Jung-ah Choi, Sang-Seok Oh, Patricia M Nuijten, Manki Song, Hyo-Young Chung, Carel A van Baalen, Ki-Sung Kwon, Soo-Young Lee

**Affiliations:** Biotechnology Research Institute, Celltrion Inc., Incheon, Republic of Korea; Science Unit, International Vaccine Institute, Seoul, Republic of Korea; New Drug Development Center, Osong Medical Innovation Foundation, Cheongju, Republic of Korea; Viroclinics Biosciences, Rotterdam, The Netherlands

**Keywords:** SARS-CoV-2 virus, P.1, Brazil variant, CT-P59, Regdanvimab, Therapeutic antibody

## Abstract

P.1. or gamma variant also known as the Brazil variant, is one of the variants of concern (VOC) which appears to have high transmissibility and mortality. To explore the potency of the CT-P59 monoclonal antibody against P.1 variant, we tried to conduct binding affinity, *in vitro* neutralization, and *in vivo* animal tests. In *in vitro* assays revealed that CT-P59 is able to neutralize P.1 variant in spite of reduction in its binding affinity against a RBD (receptor binding domain) mutant protein including K417T/E484K/N501Y and neutralizing activity against P.1 pseudoviruses and live viruses. In contrast, *in vivo* hACE2 (human angiotensin-converting enzyme 2)-expressing TG (transgenic) mouse challenge experiment demonstrated that a clinically relevant or lower dosages of CT-P59 is capable of lowering viral loads in the respiratory tract and alleviates symptoms such as body weight losses and survival rates. Therefore, a clinical dosage of CT-P59 could compensate for reduced *in vitro* antiviral activity in P.1-infected mice, implying that CT-P59 has therapeutic potency for COVID-19 patients infected with P.1 variant.

**Highlights:**

- CT-P59 could bind to and neutralize P.1 variant, but CT-P59 showed reduced susceptibility in *in vitro* tests.
- The clinical dosage of CT-P59 demonstrated *in vivo* therapeutic potency against P.1 variants in hACE2-expressing mice challenge study.
- CT-P59 ameliorates their body weight loss and prevents the lethality in P.1 variant-infected mice.

## 1. Introduction

On November 2021 in Brazil, SARS-CoV-2 infection rapidly increased again in Manaus, where previously showed high sero-prevalences and the newly emerging virus was designated as P.1 or N501Y.V3 [1, 2]. Currently, the P.1 variant become a dominant variant in Brazil and transmitted in many countries [1]. Intriguingly, P.1 variants as well as Delta variant (B.1.617.2) showed rapid displacement of B.1.1.7 in the Unites States [3].

It’s known that the P.1 variant possesses 12 amino acid mutations (L18F, T20N, P26S, D138Y, R190S, K417T, E484K, N501Y, D614G, H655Y, T1027I, and V1176F) in the spike protein of SARS-CoV-2 viruses [4]. In particular, three mutations (K417T, E484K, and N501Y) in RBD (Receptor Binding Domain) of P.1 are common in B.1.1.7 (N501Y) and B.1.351 (K417N, E484K, and N501Y) which are associated with increased transmissibility, immune escape and pathogenicity [4–7]. N501Y and E484K mutations in RBD was shown to increase the binding affinity to ACE2 (Angiotensin-Converting Enzyme 2) cellular receptor without any change of spike protein expression level [8]. Moreover, E484K and K417N/K417T mutations were known to reduce COVID-19 vaccine efficacy and confer resistance to some therapeutic monoclonal antibody such as Bamlanivimab or Etesevimab [4, 7, 9, 10]. Therefore, P.1 has potential to be rapidly transmissible and resistant to monoclonal antibody therapy as well as vaccines.

The therapeutic monoclonal antibody, CT-P59 which binds to RBD of SARS-CoV-2, interfering with access of ACE2 (Angiotensin-Converting Enzyme 2) receptor for viral infection, has demonstrated that CT-P59 has high potency against the original SARS-CoV-2 in *in vitro* and *in vivo* studies with hamster, ferret, and monkey [11]. Previously, we reported that a therapeutic dose of CT-P59 which is relevant to human dosage showed *in vivo* efficacy against B.1.351 variants in ferret animal model, despite the reduction in *in vitro* neutralizing activity [12]. Although P.1 has in common with three mutations sites of concern with B.1.351, it remains elusive whether CT-P59 can neutralize P.1 variant. To address this question, we performed *in vitro* binding, neutralization assays and *in vivo* challenge experiment in hACE2-expressing mice.

## 2. Materials and methods

### 2.1. Cells and viruses

For microneutralization assay, VeroE6 cells (ATCC, CRL-1586) were grown as previously described [12] and Brazil P.1 variant (Isolate hCoV-19/Japan/TY7-503/2021) were obtained through BEI Resources (Catalog No. NR-54982). Derivatives of HEK-293T cells expressing ACE2 were generated by transducing HEK-293T (ATCC, CRL-3216) cells with ACE2 (Addgene, #145839). Cells were used as single cell clone was derived by limiting dilution from the bulk populations. HEK293T-ACE2 cells were cultured in Dulbecco’s modified Eagle’s medium (DMEM) supplemented with 10% (v/v) fetal bovine serum (FBS) and 2 mM penicillin-streptomycin (100 U/mL). K417T, E484K, N501Y, and P.1 pseudoviruses were generated by gene cloning and confirmed by sequencing and Western blot analysis. For *in vivo* mice challenge study, hCoV-19/Korea/KDCA95637/2021 (P.1) were provided by National Culture Collection for Pathogens (#43388).

### 2.2. Biolayer interferometry (BLI)

Binding affinity of CT-P59 to wild type and mutant SARS-CoV-2 RBD were evaluated by the Octet QK^e^ system (ForteBio) using a modification of a method as previously described [11]. The P.1 triple mutant SARS-CoV-2 RBD (K417T/E484K/N501Y) was purchased from Sino Biological.

### 2.3. Pseudovirus assay

Luciferase-based pseudovirus assay was carried out using wild-type and mutant spike-expressing lentiviruses. Mutations in spike gene were introduced by gene cloning method and its protein expression was confirmed by Western blot analysis. The pseudoviruses were produced by transfection with luciferase reporter plasmid along with Gag-Pol, Rev, and Spike expression plasmid and then the copy number of pseudoviruses was quantitated by qPCR. Pseudoviruses were mixed with diluted antibodies ranging either from 100 to 0.005 ng/mL or 1000 to 0.05 ng/ml. The inocula infected ACE2-expressing HEK293T cells. After 72 h, luciferase activities were measured and IC_50_ values were calculated with Prism.

### 2.5. Microneutralization (ViroSpot) assay

To examine the susceptibility of SARS-CoV-2 variants, a microneutralization assay was performed as previously described [12]. Briefly, virus was mixed with serial dilutions CT-P59 and then the virus/CT-P59 mixture was added to Vero E6 cells. After a subsequent incubation, carboxymethylcellulose (CMC) overlay was added. The assay plates were incubated for 16-24 hours. The cells were fixed and permeabilized, followed by treatment with a murine anti-nucleocapsid monoclonal antibody (Sino Biological), a secondary anti-mouse IgG peroxidase conjugate (Thermo Scientific) and TrueBlue (KPL) substrate. Images of all wells were acquired by a CTL Immunospot analyzer, equipped with Biospot^®^ software to quantitate the nucleocapsid-positive cells.

### 2.6. Animal experiments

8-week-old female human ACE2 transgenic mice, tg(K18-ACE2)2Prlmn, were purchased from The Jackson Laboratory (ME, USA). Mice were housed in a certified A/BSL-3 facility at the International Vaccine Institute (IVI). Before virus inoculation, all mice had an acclimatization period of one week. All procedures were approved by the Institutional Animal Care and Use Committee at the IVI (IACUC Approval No. 2020-021).

### 2.7. Mouse study

Mice (n=11/group) were intranasally inoculated with 1×10^4^ PFU/30 μL of P.1 variant under anesthesia. After 8 h virus challenge, four different doses (5, 20, 40, and 80 mg/kg) were administered intraperitoneally. Control mice were administered with formulation buffer via the same route. Viral load was measured from nasal wash and lung at 3 dpi or 6 dpi (days post infection), and 4 animals were euthanized at each day for virus quantification. Mortality was recorded from 7 dpi to 10 dpi from three animals per group after final scheduled euthanasia, and body weight of all remaining mice were monitored daily throughout the study period. Animals showing more than 30% loss of body weight were euthanized and excluded from statistical analysis.

### 2.8. Virus titration and quantitation

To assess SARS-CoV-2 viral loads in the challenged transgenic mice, lung tissues and nasal washes were collected. The lungs from each mouse were aseptically removed and washed with Hank’s balanced salt solution containing 1% penicillin/streptomycin (Thermo Fisher Scientific). The nasal washes from each mouse were collected by flushing with 50 μL of sterilized PBS twice through the nasal cavity and stored at −80°C until use. The lung tissues were homogenized and strained through a 70-μm cell strainer (Becton Dickinson). The collected supernatants were stored at −80°C. The lung samples and nasal washes were 4-fold serially diluted in DMEM and inoculated to Vero E6 cells in a 12-well plate. After incubation for 30 min, the inoculum was removed and covered with agar-overlay media including 1% (w/v) low-melting agarose (Lonza) and 2% FBS in DMEM. Following incubation at 37°C for 3 days, the cells were fixed with 4% (v/v) formaldehyde, and stained with 0.05% crystal violet (Sigma). The number of plaques per well was counted and analyzed.

## 3. Results

### 3.1. *In vitro* binding affinity and susceptibility tests

To investigate the therapeutic efficacy of CT-P59 against P.1 variant, we first determined the binding affinity of CT-P59 against triple mutant RBD (K417T/E484K/N501Y) by using BLI (Bio-Layer interferometry). The equilibrium dissociation constant (K_D_) of CT-P59 against the P.1 triple mutant RBD was reduced by approx. 12-fold compared to that against wild-type RBD (Table 1 and Supplementary Fig. 1). Next, to assess the susceptibility of P.1 variant to CT-P59, we conducted two sorts of cell-based assays with live viruses or pseudotyped viruses. In SARS-CoV-2 spike expressing pseudovirus assay, CT-P59 significantly inhibited SARS-CoV-2 D614G pseudovirus with IC_50_ value of 0.219 ng/mL, but showed approx. 61-fold reduced neutralization of P.1 pseudotyped viruses with an IC_50_ value of 13.45 ng/mL (Table 2 and Supplementary Fig. 2A). While CT-P59 showed lower IC_50_ value against K417T mutant pseudovirus than that against D614G, E484K and N501Y was less than 10-fold susceptible to CT-P59, which showed approx. 8.7-fold and 5.5-fold reductions. In addition, the live virus neutralization assay showed approx. 138-fold reduced susceptibility against P.1 variants, compared to that against wild type SARS-CoV-2 (Table 2 and Supplementary Fig. 2B). Thus, we found that CT-P59 is able to neutralize P.1 variants but showed reduced binding and antiviral ability in *in vitro* experiments.

**Table 1.** Binding affinity between CT-P59 and P.1 mutant RBD.

| Ligand<sup>1</sup> | Analyte | $k_{on}$ ($M^{-1}s^{-1}$) | $k_{off}$ ($s^{-1}$) | $K_D$ (M) | Fold change |
| --- | --- | --- | --- | --- | --- |
| RBD (wild-type) | CT-P59 | 9.32E+05 | 5.89E-05 | 6.32E-11 | - |
| K417T/E484K/N501Y | CT-P59 | 6.80E+05 | 5.23E-04 | 7.69E-10 | 12 (reduction) |

<sup>1</sup>RBD (wild-type) was prepared in-house and P.1 triple mutant RBD was purchased from Sino Biological.

**Table 2.** Neutralization effect of CT-P59 against P.1 live viruses and P.1 pseudoviruses.

| Viruses | IC<sub>50</sub> (ng/mL) | Fold change (reduction) |
| --- | --- | --- |
| P.1 (live virus) | 275.75 (WT: 2.0) | 137.86 |
| P.1 (Pseudovirus) | 13.45 (D614G: 0.219) | 61.42 |
| K417T | 0.154 (D614G: 0.219) | 0.70 |
| E484K | 3.273 (D614: 0.378) | 8.66 |
| N501Y | 1.202 (D614G: 0.219) | 5.49 |

### 3.2. *In vivo* efficacy of clinical dosages of CT-P59 in mouse model

To evaluate whether CT-P59 is able to reduce viral loads and protect from exacerbating the infection against P.1 variants in *in vivo* setting, mice were treated with clinically relevant dosages considering the 40 mg/kg of clinical dose of CT-P59. The dose of 80 mg/kg was selected by equivalent exposure in mice to that of clinical study (data not shown), and lower doses of 5 and 20 mg/kg were additionally selected for further investigation with the lower exposure than the clinical situation. P.1 variants severely affected the survival rate of control animals resulting in 0% survival rate at 8 dpi (days post-infection), having two mice euthanized which showed more than 30% body weight loss at 7 dpi and one mouse died at 8 dpi. In contrast, no death or over 30% loss of body weight was observed in all CT-P59 treatment groups throughout the study period (Fig. 1 A). Following infection with P.1 variants, body weights were significantly not altered in CT-P59 treatment groups. In contrast, the control groups infected by P.1 only lose the body weight since 1 dpi, resulting in severely decreased mean body weights and eventually 28.4% loss at 7 dpi. Importantly, the CT-P59 treatment delayed statistically significant loss of their body weight compared to control mice from 2 dpi to 6 dpi. CT-P59 treatment groups at 5, 20, 40 and 80 mg/kg showed 18.8, 16.6, 16.7 and 9.2% loss of mean body weight at 6 dpi, respectively. However, the mean body weight of 20, 40 and 80 mg/kg CT-P59 treatment groups began to revitalize from 6 dpi and fully recovered at 10 dpi (Fig. 1 B). To monitor viral loads in upper and lower respiratory tracts, we performed plaque assay with mice lung and nasal wash samples. CT-P59 treatment effectively reduced the infectious virus titers in lung tissues and nasal washes of the mice challenged with P.1 variants. The mean virus titer in lung tissues reached 5.1 log (PFU+1)/mL at 3 dpi and declined to 2.4 log (PFU+1)/mL at 6 dpi in the control group. In contrast, all CT-P59 animals had reduced viral titers at 3 dpi and 6 dpi. At 3 dpi, infectious viral titers were 9.2, 5.5, 2.5 and 3.6-fold log reduced in 5, 20, 40 and 80 mg/kg CT-P59 treatment groups respectively, compared to the control group; complete diminution were observed in lungs from 3 out of 4 animals in 5, 20 and 80 mg/kg treatment groups. At 6 dpi, no viral titers were detected in all CT-P59 treatment groups showing a complete reduction of infectious viruses (Fig 1 C and D). In upper respiratory tracts, the mean virus titers from nasal washes reached 0.5 log (PFU+1)/mL at 3 dpi and showed a value of 0.6 log (PFU+1)/mL at 6 dpi in the control group; one mouse from control group showed viral loads of 2 log (PFU+1)/mL or above at 3 dpi and 6 dpi, respectively. On the contrary, no infectious viruses were measured in all CT-P59 treated groups at 3 dpi and 6 dpi, in line with lower respiratory tracts as stated above (Fig. 1 E and F).

**Figure 1.**
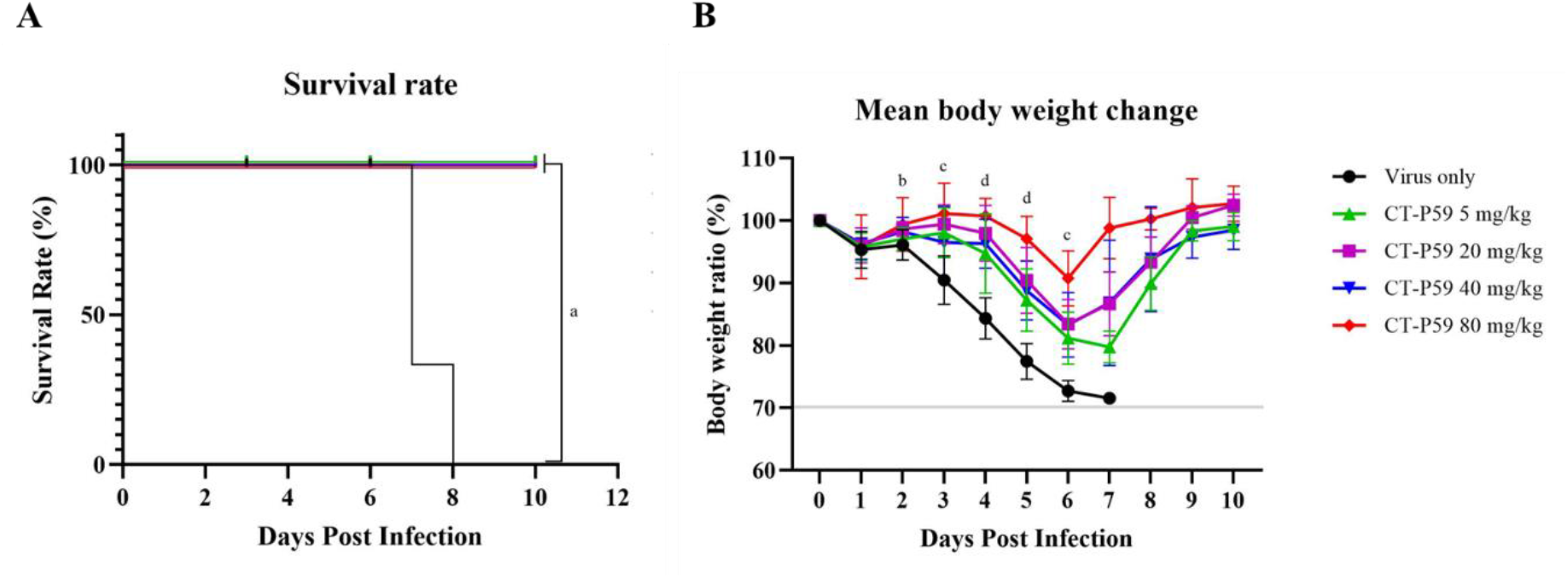
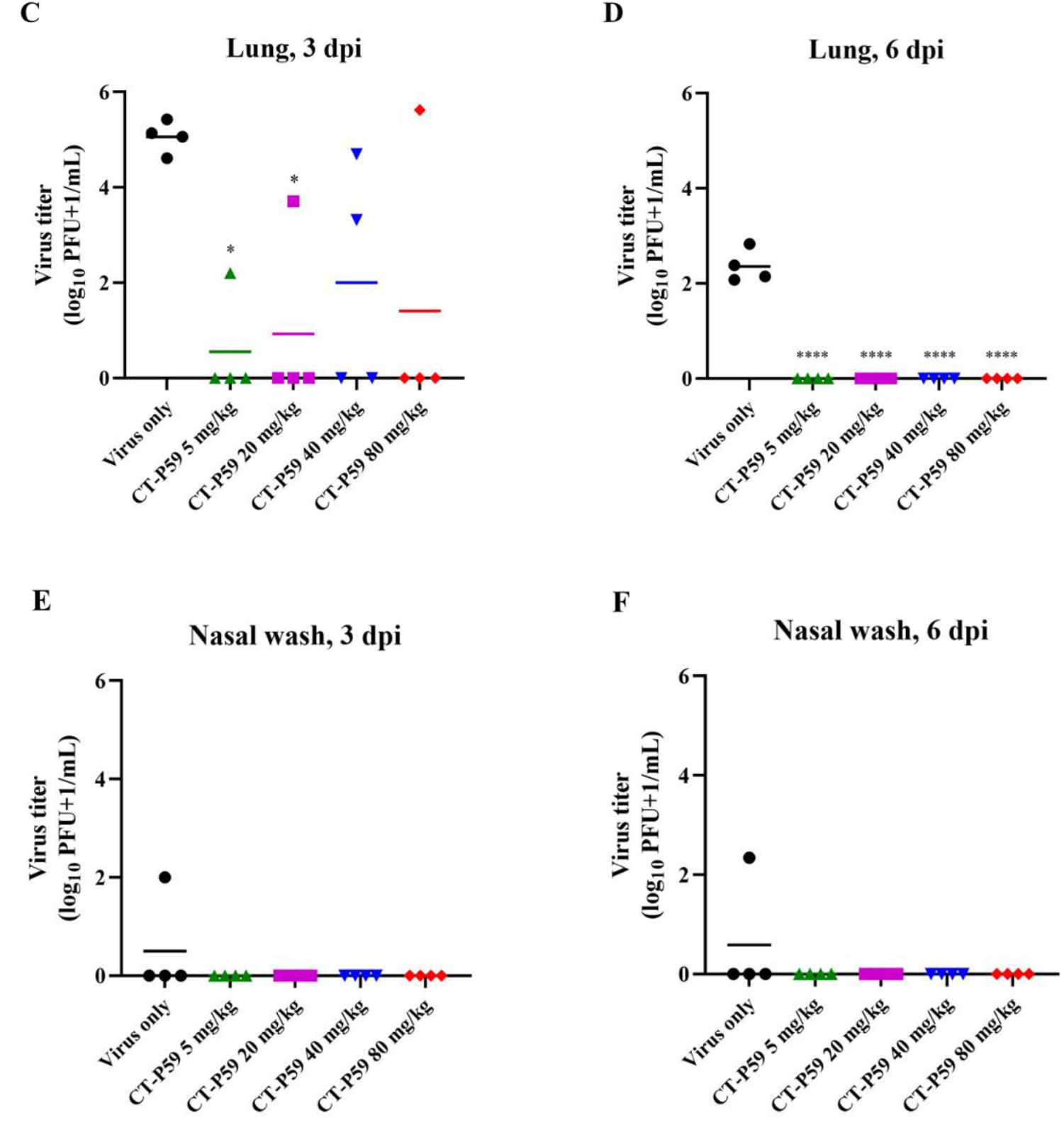
*In vivo* efficacy of CT-P59 against P.1 variant in transgenic mouse model. Human ACE2 Transgenic mice (n=11/group) were challenged with 10^4^ PFU of P.1 variants. Vehicle and 5, 20, 40 and 80 mg/kg of CT-P59 were administered intraperitoneally 8 h after virus inoculation. Body weights and survival rates were monitored daily until 10 dpi (A, B). 30% or higher of weight loss was considered to be dead. Four animals each were euthanized for virus titration at 3 dpi and 6 dpi. The virus titers from lung tissue and nasal wash were measured (C, D, E, F) using plaque assay. Alphabets and asterisks indicates statistical significance between the control and each groups as determined by one-way ANOVA followed by a Dunnett’s post-hos test. *a* denotes p<0.0001 between control and CT-P59 treatment groups (pooled). *b* denotes p<0.01 to p<0.05 between control and CT-P59 treatment group at 80 mg/kg. *c* denotes p<0.0001 to p<0.01 between control and CT-P59 treatment groups at 5, 20, 40 and 80 mg/kg. *d* denotes p<0.0001 to p<0.001 between control and CT-P59 treatment groups at 5, 20, 40 and 80 mg/kg. * indicates *P* < 0.05, and **** indicates *P* < 0.0001.

## 4. Discussion

In this study, as with previous report on B.1.351 [12], we demonstrated that the therapeutic effect of CT-P59 against the P.1 variant of SARS-CoV-2 by *in vitro* neutralization tests and *in vivo* challenge experiments using hACE2 transgenic mice. Although P.1 share common mutation locations at K417, E484, and N501, the neutralization against P.1 was observed to differ from that on B.1.351 as mention above, since CT-P59 showed 61-fold and 138-fold reduced neutralizing activity against P.1 with the pseudovirus assay and micro-neutralization assay, respectively. Therefore, substitution or deletion mutations in the other regions such as NTD (N-terminal domain) or S2 except RBD in P.1 or B.1.351 variant could contribute to change in conformation of spike protein, potentially impacting the neutralization of therapeutic antibodies.

In previous report, B.1.351 known as South African variant or beta variant showed 20-fold and 33-fold reduced susceptibility to CT-P59 by micro-neutralization assay and pseudovirus assay, respectively. However, a therapeutic dosage of CT-P59 which corresponds to clinical dosage for patient treatment has inhibitory effect on replication of B.1.351 in the ferret challenge model to a similar extent to replication of wild type virus. Afterwards, we found that CT-P59 showed 1.8-fold stronger antiviral effect on B.1.1.7 variants than wild type and 310-fold reduced neutralizing activity against B.1.351 isolated from patients in South Korea in plaque reduction neutralization test, but we confirmed the neutralizing effect of clinically relevant and lower doses of CT-P59 in B.1.351-infected hACE2-expressing mice (unpublished results). Along with P.1 data, we speculate that a clinical dosage of CT-P59 (40 mg/kg in human patients) would be able to overcome 138 ~ 310-fold reduction in *in vitro* antiviral activity.

In parallel, we determined the *in vivo* neutralizing potency of CT-P59 against P.1 variant in the hACE2 TG mouse model. K18-hACE TG mice model was known to be susceptible to SARS-CoV-2 and develop severe clinical symptoms of COVID-19 disease [13, 14]. Although ferrets infected by wild type or B.1.351 showed lower viral titers in the lower airway compared to the upper airway [11], we found that the virus titers of P.1 variants remained lower in the upper respiratory tract than in the lower respiratory tract in the mice. These results were not surprising because K18-hACE TG mice are highly susceptible to SARS-CoV-2 infection and experience higher viral titers in lung tissues compared to the upper respiratory tract [15]. In line with this finding, a recent report showed that P.1 variant-infected common laboratory mice resulted in high viral titers in lung tissues [16]. However, neither body weight loss nor lethality was observed in wild mice challenged with P.1 variants on the contrary to our results which manifest a severe body weight loss in the control group. Given that unmodified animals infected with P.1 variants showed anatomical and histopathological changes in the lower respiratory tract [17], our findings that CT-P59 treatment led to complete reduction of infectious viruses, full recovery of body weight loss, and no death rates provide us with expectation to alleviate additional pneumonia as well as virus infections, although no pathological data were available in this study.

## Conclusion

The *in vitro* and *in vivo* studies demonstrate that clinical dosages of CT-P59 which corresponds to human dose for patients and lower doses, showed significant reduction in viral loads and protection from worsening of symptoms in P.1-infected mice despite the reduced *in vitro* potency of CT-P59 against P.1 variants.

## Acknowledgements

This research did not receive any specific grant from funding agencies in the public, commercial, or not-for-profit sectors.

## Supplementary Material

**Supplementary figure 1.**
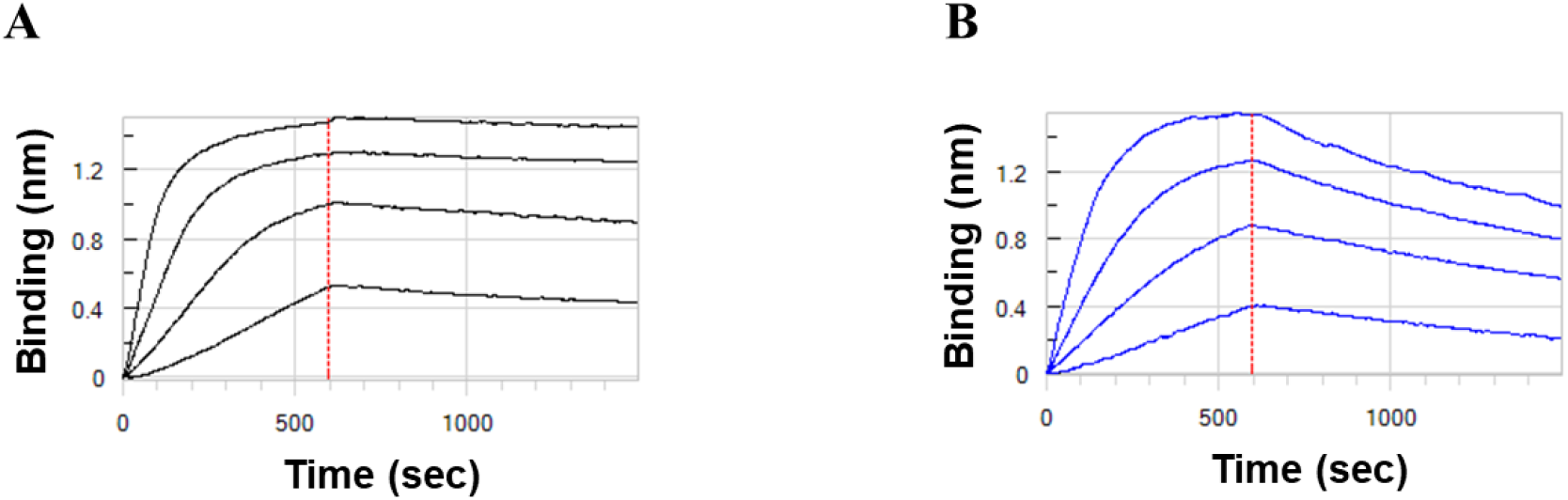
BLI sensorgrams for the evaluation of CT-P59 binding affinity against wild-type RBD and P.1 triple mutant RBD. SARS-CoV-2 wild-type RBD (A) or P.1 triple mutant RBD (B) was loaded and immobilized onto Anti-Penta-His (HIS1K) Biosensor (Sartorius) at the concentration of 50 nM for 7.5 min, and then CT-P59 was flowed with the concentration of 10 nM, 5 nM, 2.5 nM and 1.25 nM for 10 min and 15 min to generate association and dissociation curve, respectively. BLI experiment was operated using Octet Data Acquisition v11.0 software (ForteBio), and calculation of binding affinity was performed with ForteBio Data Analysis v11.0.

**Supplementary figure 2.**
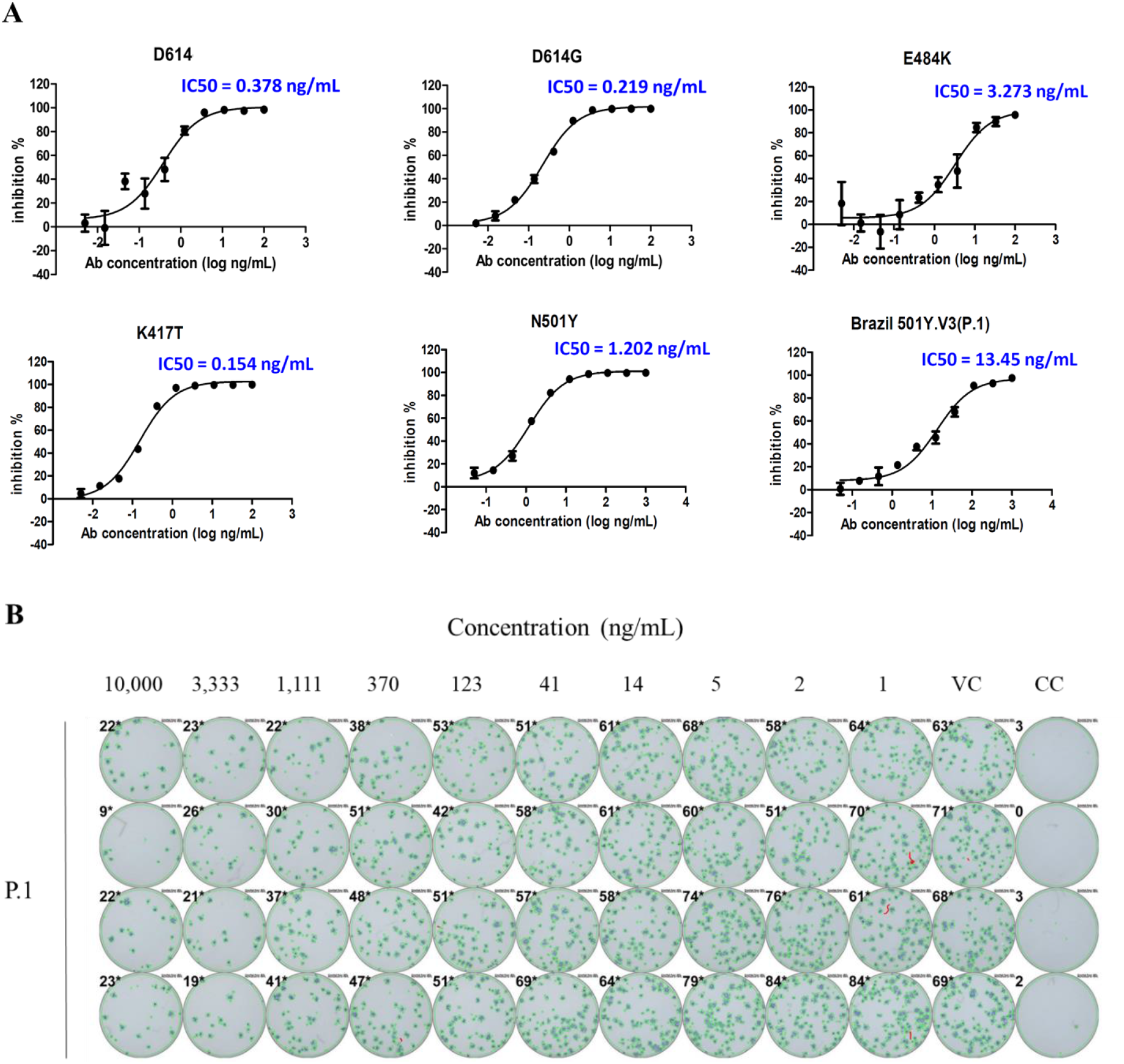
*In vitro* neutralization of CT-P59 against P.1. Serially diluted CT-P59 was mixed with indicated variant pseudoviruses for inoculation to ACE2-expressing HEK293T cells. Luciferase activity was measured and %Neutralization was calculated (A). CT-P59 dilutions were pre-incubated with P.1 variant. The antibody-virus mixture were inoculated into Vero E6 cells, incubated and probed by anti-nucleocapsid antibody and staining (B). VC and CC represent as virus control and cell control, respectively.

